# Stereotactic system for accurately targeting deep brain structures in awake head-fixed mice

**DOI:** 10.1101/539957

**Authors:** Yonatan Katz, Michael Sokoletsky, Ilan Lampl

## Abstract

Deep brain nuclei, such as the amygdala, nucleus basalis, and locus coeruleus, play a crucial role in cognition and behavior. Nonetheless, acutely recording electrical activity from these structures in head-fixed awake rodents has been very challenging due to the fact that head-fixed preparations are not designed for stereotactic accuracy. We overcome this issue by designing the DeepTarget, a system for stereotactic head-fixation and recording, which allows for accurately directing recording electrodes or other probes into any desired location in the brain. We then validated it by performing intracellular recordings from optogenetically-tagged amygdalar neurons followed by histological reconstruction, which revealed that it is accurate and precise to within ∼100 μm. Moreover, in another group of mice we were able to target both the mammillothalamic tract and subthalamic nucleus. This approach can be adapted to any type of extracellular electrode, fiber optic or other probe in cases where high accuracy is needed in awake, head-fixed rodents.

**Highlights:**

> The Deep Target, new system for accurately targeting deep nuclei in head-fixed animals for electrophysiology and optogenetics.
> Accurate and precise to within 100 μm following a one-time alignment.
> Validation: Opto-tagged Vm recordings in the amygdala of awake mice.
> Validation: Targeting multiple deep brain structures in the same mouse.

## Introduction

In recent years electrophysiological recordings from awake animals have become standard in neuroscience research, enabling us to study circuits of perception, learning, behavior and other cognitive functions at high temporal resolution. Such recordings have been greatly aided by the development of head-fixed preparations (Chang et al., 2016; Madularu et al., 2017; Osborne and Dudman, 2014; Suga et al., 1978), wherein a headbar is fixed onto the animals’ skull to allow probes easy access to neural tissue. Compared to fixing the animal within a stereotactic device (e.g. using earbars), fixation using a headbar is less painful and distressing for animals. In order to maintain stereotactic accuracy, experimenters typically use a stereotactic device during the surgery where the headbar is implanted. However, its use during such procedures has been limited to marking the positions of the craniotomy, while the headbar itself is implanted manually by visual guidance. This may be sufficient for targeting the recording to the cortex or large brain structures in close proximity, such as the hippocampus (Bittner et al., 2017; Cohen et al., 2017; Hulse et al., 2016; Stempel et al., 2016), striatum (Ketzef et al., 2017), and thalamus (Urbain et al., 2015), but not for targeting smaller and deeper structures.

This issue may be overcome when using extracellular probes, where the multitude of unit responses can provide an indication of whether the correct area was reached. This is the case, for example, when attempting to record from sensory subcortical neurons which are driven by external stimulation (Cohen-Kashi Malina et al., 2016; Hei et al., 2014) or when using optogenetic tagging (Kravitz et al., 2013; Meir et al., 2018; Xu et al., 2015). However, difficulties are compounded for single cell electrophysiology (cell-attached and intracellular), where correct placement is more difficult to verify and the pipette also needs to be replaced and then retargeted after every attempt, causing additional damage to the brain. Moreover, time is of the essence in such experiments, as the awake animal can be head-fixed for only a limited duration. Therefore it is crucial to target the pipette to the right location from the first attempt.

For these reasons, we sought to develop a system which would allow for accurately targeting deep and small structures in head-fixed mice. Such a system would need to accomplish two goals: first, the headbars for head-fixation should be mounted in a stereotactically-accurate manner, and second, that this accuracy carry over to the recording stage. We accomplished the first goal by designing custom headbars that can be mounted using the same adapter that was used to stereotactically align the skull, and the second by placing the recording apparatus on the same plane as the head-fixation apparatus. We verified the accuracy and the precision of our approach histologically by targeting right and left amygdalas and in other mice both the mammillothalamictract and the subthalamic nucleus. We also demonstrated the stability of the system by performing intracellular recordings from optogenetically-identified amygdalar neurons in awake mice.

## Results

Accurate recordings in deep brain structures were achieved in our system in two steps. In the first, the headbar was mounted in a manner aligned with the brain’s stereotactic coordinates; in other words, it was mounted while constrained in both position (centered at bregma) and rotation (parallel to the horizontal plane). In the second step, the manipulator and thus the electrodes’ coordinates were aligned with the headbar, achieving a recording system that is stereotactically aligned to the brain. Note that in our experiments the mounting and recording steps were performed in two different experimental sets.

### Creating a stereotactically-compatible headbar

The headbar that we created is a modified version of a design originally in Osborne and Dudman (2014). It is molded from plastic in a 3D printer (ProJet 3500 HDMax, 3D Systems) and weights about 0.3 grams. Its bottom side is concave and matches the curvature of the mouse skull (diameter = 16.0 mm), creating an arc covering about a quarter of a circle (Fig. 1A,B). It also has a 2×2 mm square holding post which allows it to be attached to the stereotactic manipulator arm. Using the 3D printer we were able to produce different types of headbars, each containing an opening for a specific recording area (S1, V1, PFC, amygdala, etc.).

**Figure 1.**
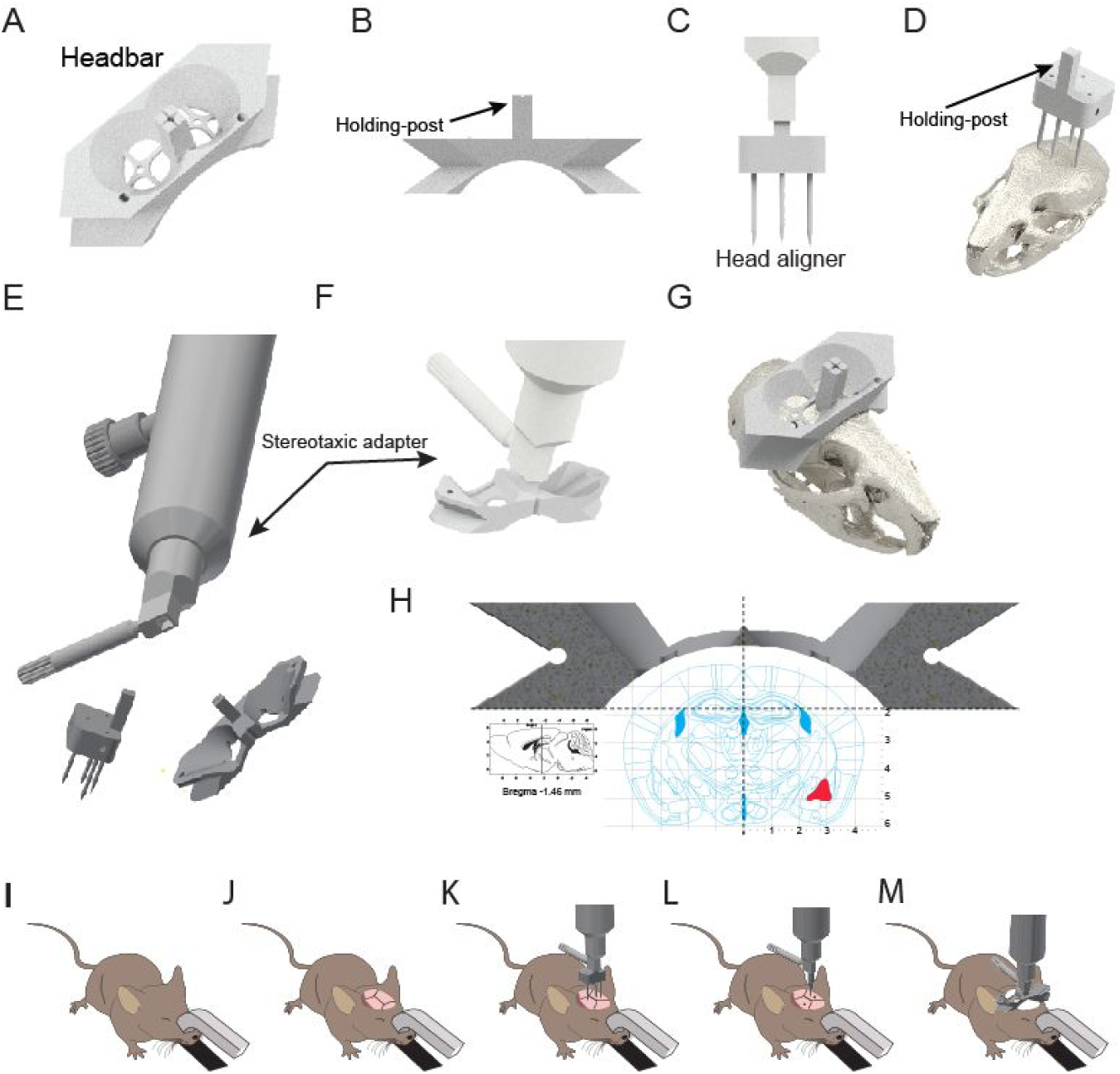
Stereotactic headbar mounting method. **A.** Isometric view of the headbar showing the target shapes (at the center of the holes) used to mark the position of the amygdala. **B.** Front view of the headbar showing the post used to attach the headbar to the stereotactic adapter, and the concave shape of its bottom which matches the curvature of the skull. **C.** Front view of the head-aligner attached to the stereotactic adapter. The pins are used for initial stereotactic alignment of the animal. **D.** Isometric view of the head-aligner and its position relative to the skull during alignment. The holding post is inserted into the stereotactic adapter at the same position as the headbar. **E.** Bottom view of the stereotactic adapter unveiling the socket in which the head-aligner and the headbar posts are inserted into. **F.** The headbar position when it is attached to the stereotactic adapter before is it mounted over the skull. **G.** The headbar mounted over the skull. **H.** A scheme of a coronal cut at −1.46 AP showing the position of the mounted headbar. The target recording location at the right amygdala is marked with red. **I-M.** A procedural scheme of the headbar mounting procedure. **I.** The anesthetized mouse is mounted at the head holder. **J.** The skull is exposed. **K.** The head is aligned in stereotactic coordinates using the head-aligner. **L.** The projected recording locations are marked on the skull following validation of stereotactic alignment. **M.** The headbar is mounted.

### Mounting the headbar

To align the skull of the animal, which is held on an articulated arm with a head-holder (Haidarliu, 1996; Slotnick, 1972), we used a custom head-aligner which consisted of four pins (Fig. 1C,D) - two pins matching the bregma and lambda position (3.8 mm apart) aligning the anteroposterior (AP) axis and two pins positioned 3 mm laterally from the midline to align the mediolateral plane (Fig. 1C). Note that the vertical distance between the ending points of the two pairs of pins matches the curvature of the skull. The head-aligner is first attached to the stereotactic manipulator arm (Fig. 1E) and then vertically lowered so to bring all four pins in close contact with the skull (Fig. 1D). If the pins were not all in close contact, we would adjust the angle of the skull in one or more of the three rotational axes and repeat the procedure until close contact was achieved. Note that although we used the head-aligner due to its speed, it is also sufficient to use a single stereotactic attachment to measure the height at the four points and then adjust accordingly, as long as the final position of the attachment is above bregma. Indeed, we performed this procedure as an additional validation of alignment.

At this stage the skull was stereotactically-aligned and we could replace the head-aligner with the headbar on the manipulator arm (Fig. 1E). The headbar was then lowered onto the skull until it was tightly positioned (Fig. 1F,G), and we confirmed alignment by observing that the printed crosshair targets of the headbar matched the marked craniotomy locations (Fig. 1H). Once the headbar was in the right position, it was lifted and a luting cement (RelyX, 3M) was applied over its concave side, before again lowering it onto the animal’s skull to attach it.

In summary, the mounting process (Fig. 1I-M) begins with fixing the anesthetized animal at the head-holder (Fig. 1I) and exposing the skull (Fig. 1J), then aligning the head using the head-aligner (Fig. 1K), followed by validating the alignment and stereotaxic marking of the craniotomy positions (Fig. 1L), and finally gluing the headbar to the skull (Fig. 1M).

### Aligning the recording pipette

We used a motorized manipulator (MX7600, Siskiyou) that could be advanced to any position on the XY plane. Generally, however, any manipulator that allows advancing the pipette vertically can be used instead.

We built a custom linear treadmill accompanied by V-shaped clamps that lock the headbar in the XY plane (Fig. 2A). The clamps fixate the headbar at the exact same position so the manipulator will be aligned with the brain regardless of the particular headbar used (i.e., headbars designed for different recording sites). Our experience with head-fixed mice has shown that when their body is slightly tilted (head-up, 20° tilt) they locomote and whisk more. Since the manipulator would also have to be tilted to maintain alignment, the entire treadmill apparatus was placed on an adjustable angle plate. Two metal wings were screwed to the treadmill (Fig. 2B) allowing the magnetic base for one or two manipulators to be mounted perpendicular to the headbar clamps (Fig. 2C).

**Figure 2.**
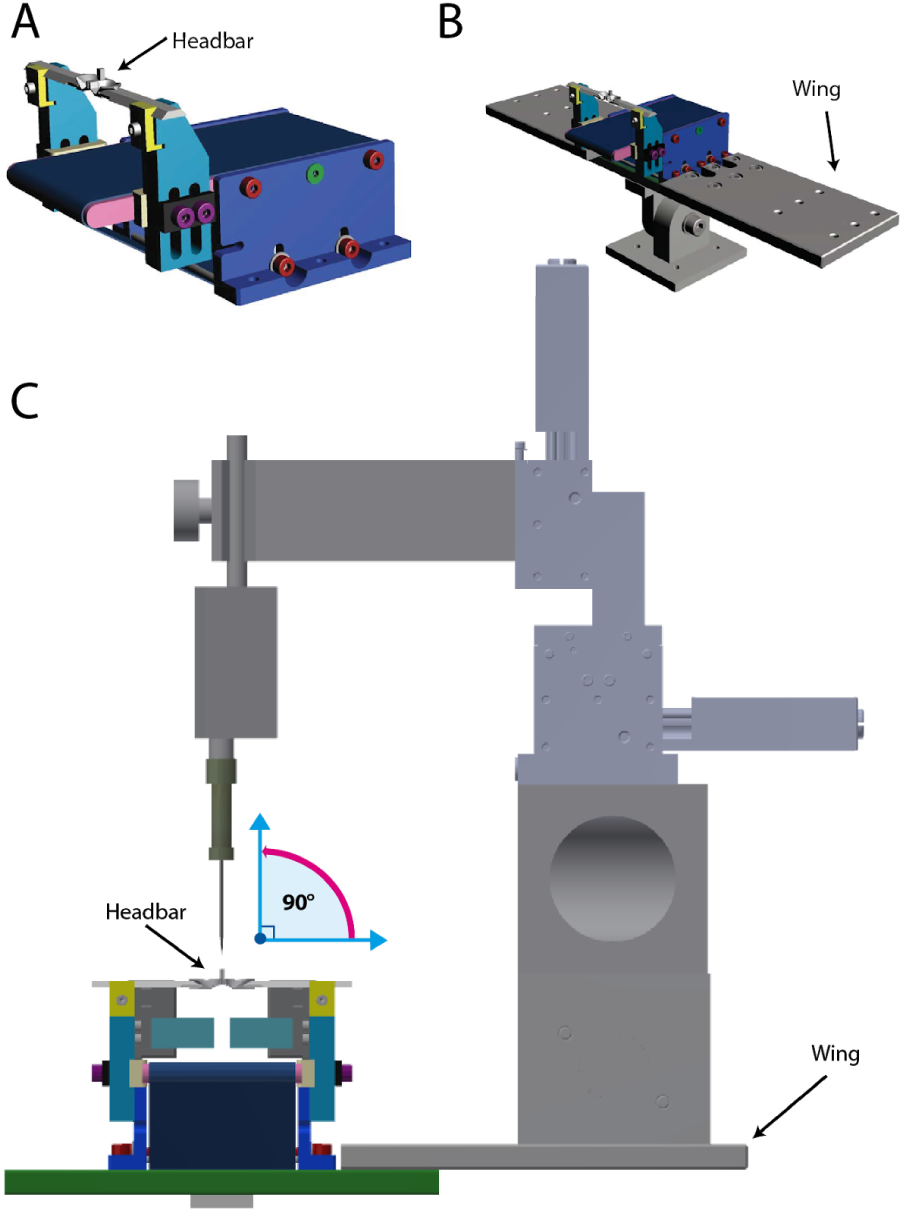
The DeepTarget system. **A.** The horizontal treadmill in which mice are mounted using the clamps that hold the headbar. **B.** The treadmill together with the side wings which hold the recording manipulator. The wings are parallel to the clamped headbar. The treadmill is sitting over adjustable angle mounting plate to position the animal at an incline to increase walking **C.** Front view of the recording manipulator mounted on the wing holding the recording pipette perpendicular to the headbar. Before recording takes place the position of the pipette is adjusted to be perpendicular both to the rostrocaudal and mediolateral planes.

### Verification of targeting with histology and extracellular recordings

To verify the accuracy and precision of the system in targeting the recording electrode to deep brain structures, we used the transgenic mouse line Thy1-ChR2-YFP which has a distinctive YFP expression in the amygdala (Fig. 3B). The amygdala of the mice is ∼0.5 mm wide when measured in the mediolateral axis and is located about 4 mm deep below the pia. Thus, high accuracy is needed to target amygalar cells. The positioning was tested using both ChR2-assisted recordings and histologically by inserting a DiI-coated pipette following the recordings. To perform the first test, we used the optopatcher (Katz et al. 2013), which enabled us to perform cell-attached recordings with light illumination deep below the cortex. Cell-attached recordings from the left (Fig. 3A) and right (Fig. 3C) hemispheres showed that cells in the targeted region reliably fired shortly after light onset (3-5 ms) and exhibited a very small jitter (10 trials superimposed), indicating that they expressed ChR2. Histology confirmed that the amygdala was successfully targeted (Fig. 3B). Another example from a different animal showed responses with similar latencies from both hemispheres and similar positioning in the histological reconstruction (Fig. 3D-F). Overall, we bilaterally targeted the amygdala in 6 animals and were able to identify 18 neurons that responded with short latency in all of them (<10 ms). We also quantified both the accuracy (absolute distance from targeted coordinate) and precision (distance from the mean) of the DiI markings in the mediolateral (ML) dimension (Fig. 3G), finding a mean accuracy of 103 ± 22 μm and precision of 77 ± 16 μm.

**Figure 3.**
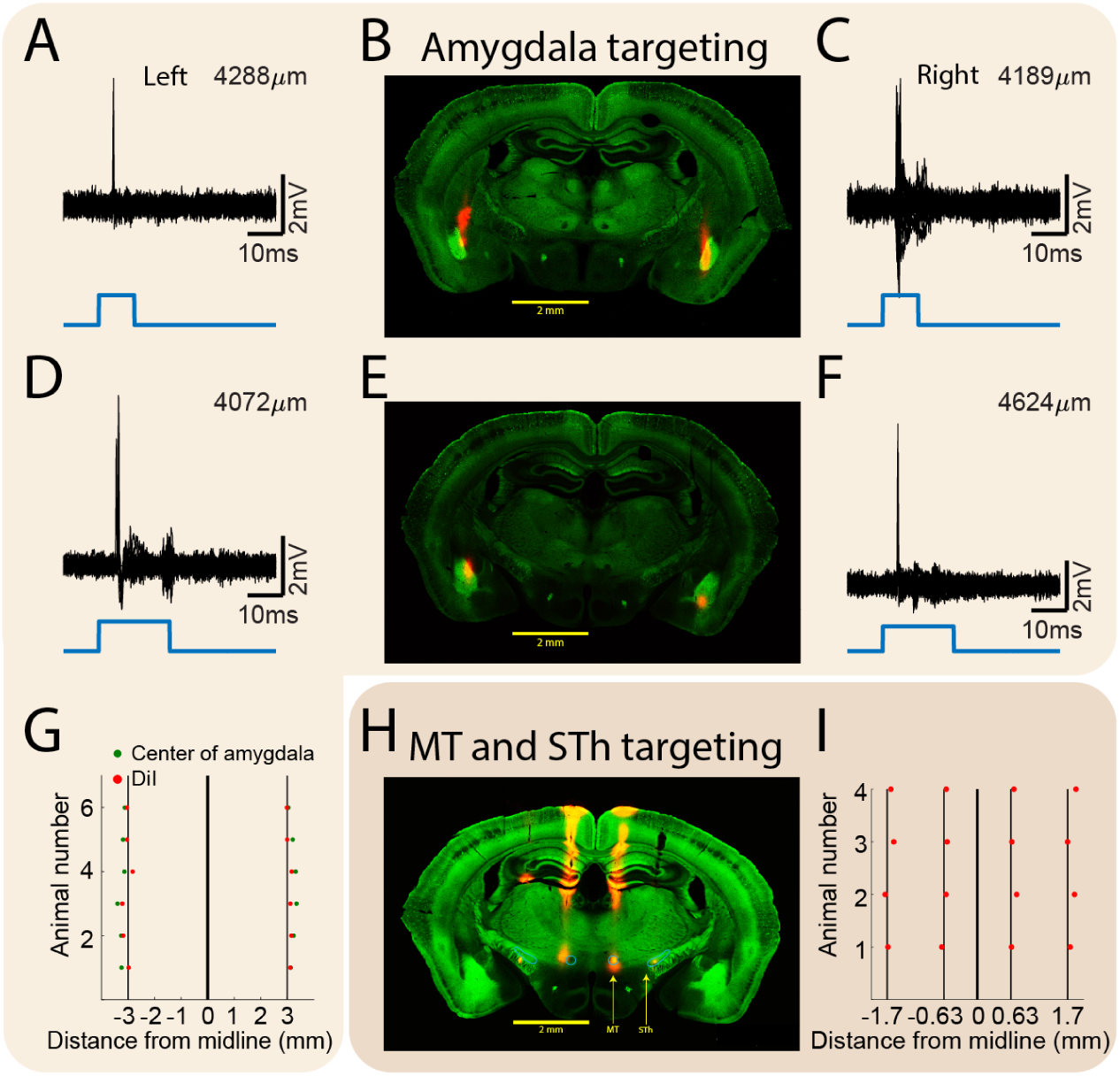
Physiological and histological verification of the DeepTarget positioning. **A.** Ten superimposed trials of responses to light stimulation of single units recorded juxtacellularly on the left hemisphere. **B.** Histological verification of the recording position of A and C using DiI marking. The amygdalar complex endogenously expresses YFP at this transgenic line and appears as a green ellipsoid on both sides. **C.** Same as A for a unit recorded on the right side. **D-F.** Same as A-C, from another animal. **G.** ML location of the center of the amygdala (green) and DiI marks (red) in both hemispheres relative to the midline (n = 6 animals), while the black lines mark the targeted coordinates. **H.** Example for DiI histology of a coronal slice of 4 different locations marked using the DeepTarget in-vivo (mammilothalamic tract (MT) and the subthalamic nucleus (STh)). **I.** ML location of the DiI marks (red) directed to the MT and STh in both hemispheres relative to the midline (n = 4 animals).

In order to further test the accuracy of the system, we targeted two additional notable and relatively small structures in both hemispheres – the mammillothalamic tract (MT) and subthalamic nucleus (STh) in 4 animals. For that we used a different headbar which enabled us to perform more medial craniotomies. We directed the electrode at the same coronal plane (−2.0 mm AP from bregma) to the MT (0.63 mm ML, 4.6 mm DV) and the STh (1.7 mm ML, 4.45 mm DV). Histology confirmed that these structures were successfully targeted (Fig. 3H), with a mean accuracy of 48 ± 11 μm and precision of 34 ± 13 μm in the ML dimension (Fig. 3I). Overall, we validated that at least three different regions in the brain can be targeted using the system with a single attempt, minimizing brain damage as well as experiment time.

### Intracellular amygdalar recordings in awake animals

As an example of an application of the system we created, we recorded the membrane potential in an amygdalar cell of an awake Thy1-ChR2-YFP mouse using the optopatcher (Katz et al., 2013). During this recording we monitored various behavioral parameters such as pupil size, whisking and locomotion, which are typically associated with arousal state. In contrast to the large ongoing activity in the cortex during quiescent periods (Meir et al., 2018; Pinto et al., 2013; Polack et al., 2013), we found that subthreshold ongoing activity in the amygdala was relatively low (Fig. 4A). During arousal states, marked by increased pupil size, whisking and locomotion activity, subthreshold activity became more noisy. In order to validate that the recording was made from an amygdalar cell we stimulated it with 7 Hz light pulses (Fig.4B). The cell showed a strong and reproducible response to light and demonstrated negligible jitter in firing (n = 10 trace, Fig. 4C). The recording position within the amygdala was also validated as before by the insertion of a DiI-coated pipette at the same position following the intracellular recording (Fig. 4D).

**Figure 4.**
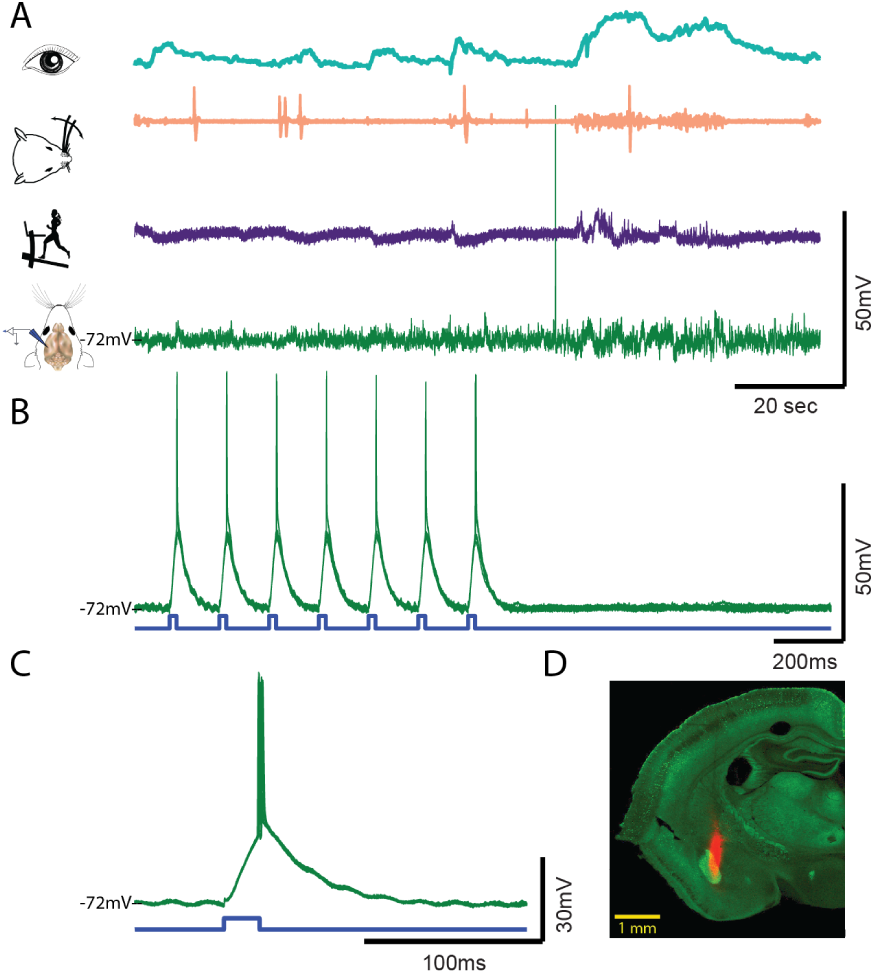
Patch-clamp amygdalar recording in an awake mouse. **A.** A two minute recording of intracellular membrane potential in the amygdala (green trace) together with various behavioral parameter such as the pupil size (cian trace), whisking (peach trace) and locomotion activity (purple trace). **B.** Response of the same cell to a 7 Hz light stimulation (n = 10 traces). **C.** Response to a single light stimulation with an expanded time scale (n = 10 traces). **D.** Histological reconstruction of the recording position using a DiI marking.

## Discussion

The head-fixed mouse preparation has been developed over a decade ago (Boyden and Raymond, 2003), and since then has been used in a wide variety of studies examining cortical activity in awake animals. Intracellular electrophysiology in such preparations has been used to uncover the impact of brain state on cortical dynamics (Poulet and Petersen, 2008), the role of inhibition on sensory response (Haider et al., 2013; Yu et al., 2016), the underlying synaptic mechanisms of feature selectivity (Cardin et al., 2007), the membrane potential correlates of detection of perception (Sachidhanandam et al., 2013), and the underlying mechanisms of active sensation (Pluta et al., 2017; Urbain et al., 2015). Clearly, to better understand cognitive and behavioral functions we must reveal the subthreshold dynamics of neurons and find how they integrate their synaptic inputs in the awake state. Nonetheless, only a small number of such studies involved diving beneath the cortical sheet (Bittner et al., 2017; Cohen et al., 2017; Hulse et al., 2016; Ketzef et al., 2017; Urbain et al., 2015), all directed to large structures. Perhaps the major reason for this is the difficulty in targeting the recording electrode into specific deep brain regions.

In this study we introduce the DeepTarget, a stereotactic apparatus enabling the accurate and precise insertion of recording or stimulation probes into deep brain structures in head-fixed awake mice. We demonstrate the power of the system using histology and recordings of optotagged neurons in the amygdala, as well as histology in two additional brain regions. It appears that the system can be generally used to target any brain structure within mice or other head-fixed small animal. Our system’s ability to accurately target deep brain structures results from alignment between its various components: the brain, the headbar, the head-fixation clamps and the recording pipette. At each stage the XY plane (horizontal) is parallel for all the components enabling the transfer of coordinates from the brain to the recording pipette.

Although our system has shown a high (∼100 μm) accuracy, it is not perfect. Variations in the location of the staining stem from at least three sources: mechanical, experimental and biological. The mechanical source arises from imperfections in the manufacturing process causing differences between the position of the head-aligner and the headbar. This difference was minimized by marking the position of the amygdala on the skull using the stereotaxic device and making minor adjustments in the headbar position to match them if needed. The experimental source can result from issues in the alignment of the animal by the experimenter and the levelling of the manipulator versus the headbar. The biological source is the result of variations between animals (Wahlsten et al., 1975), which is why we recommend to use age-matched unisexual animals. However, such variation exists even when controlling for age and sex. In our experiments the variation in the distance of the center of the amygdala from midline, based on YFP markers, was only slightly smaller (SD = 103 μm, n = 6) than the variation in DiI deposit locations (SD = 111 μm as shown above). Due to this biological variation, any improvement in the precision and accuracy of our system may not provide additional benefits when targeting deep structures.

In summary, up until now deep and small brain structures have been out of accurate experimental reach in head-fixed animals. The DeepTarget extends our ability to uncover intracellular mechanisms at any desired region within the brain. The system can also be used for acute experiments in awake animals to accurately target extracellular electrodes, such as NeuroPixels (Jun et al., 2017) optrodes (Wang et al., 2010), fiber photometry (Gunaydin et al., 2014) and any other probe.

## Methodological Considerations

### Instrument modifications

Several modifications were made to the stereotactic manipulator arm (Micro Manipulator 1760, Kopf) for this procedure. First, a square hole (2×2 mm) custom adapter was placed at its end, in order to fit the posts on both the head-aligner and headbar (Fig. 1E). During the last steps of the alignment procedure, this hole was located directly on top of bregma regardless of what was attached to it. Another modification we made was to add to the arm a sliding cylinder that can be locked at any desired height, which allowed us to rapidly switch between the head-aligner and headbar without needing to move the manipulator arm.

### Mechanical errors

Commercial systems which are mass produced have little variations in parts, while in our case mechanical inaccuracies in the prototyping manufacturing can give rise to various mechanical errors. One such error, for example, can arise from the arrangement of the headbar-aligner, in that case if the lateral pins responsible for the roll axis are not exactly aligned to the same height the skull will be tilted. In addition, the mechanical design of some parts can be further improved in order to increase the accuracy. One example for that is the stereotactic adapter which holds the headbar or the head-aligner. Its connector has a square profile into which the headbar is inserted, this results in a small degree of freedom between them which can be solved by using a pyramidal shape adapter instead of squared profile.

### System aligning

There are minor variations between different mice lines. These differences also exhibited in brain structures. Since the goal of the DeepTarget is to target small brain nuclei, small variations in the coordinates of brain structures will impair the precision of the probe targeting. Thus the best practice will be to initially mark the position of the targeted brain structure with DiI (or other marker) and to calibrate the exact coordinates in one’s mice and one’s system. Once the system is calibrated to these animals, no changes are needed when producing headbars for other brain structures.

The preferred method to validate the probe’s position is to use a reporter line expressing an excitatory opsin at the area of interest. In that case by using a system which gives optogenetic stimulation, one can have immediate feedback that the right position was achieved. This can be done using the Optopatcher or optrodes. Although the DeepTarget is relatively accurate, by using feedback one can validate the relative position within a specific nucleus with high confidence. For example, the most upper amygdalar neurons are located at a depth of about 3.8 mm but due to the diagonal nature of the structure if optogenetic feedback during recording is obtained at a deeper depth, one can estimate how to relocate the electrode in order to specifically target the dorsal cells of the BLA.

### Key Resources Table

| REAGENT or RESOURCE | SOURCE | IDENTIFIER |
| --- | --- | --- |
| **Chemicals, Peptides, and Recombinant Proteins** |  |  |
| Dil | Thermo Fisher Scientific | Cat# D282 |
| Surgical tissue adhesive | 3M | Vetbond |
| Buprenorphine | Vetmarket | 163451 |
| Carprofen | Norbrook | Norocarp |
| Silicone elastomer | Smooth-On, Inc. | Body Double™ |

Experimental Models: Organisms/Strains

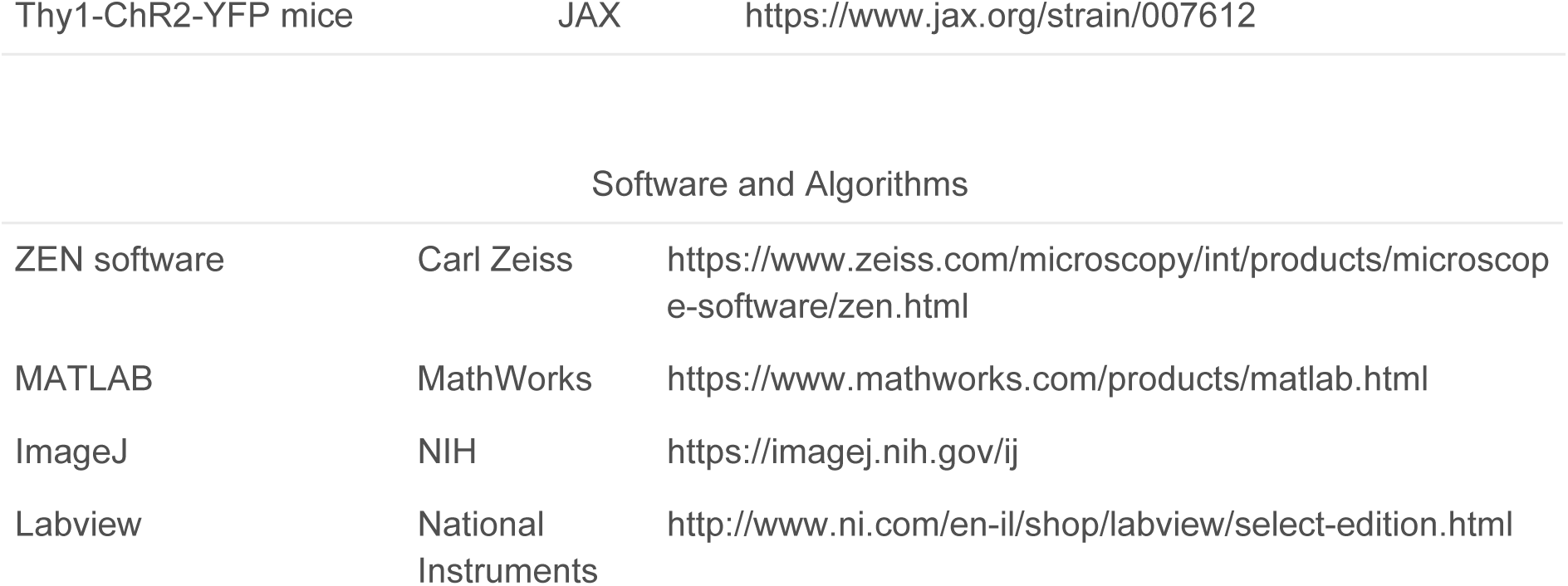

## Materials and methods

All experiments were conducted according to the Weizmann Institute Institutional Animal Care and Use Committee.

### Animals

We used ten Thy1-ChR2-YFP mice (JAX #007612, The Jackson Laboratory, RRID:IMSR_JAX:007612 (Arenkiel et al., 2007)), 8- to 15-week-old of either sex housed up to five in a cage with a 12-hr/12-hr dark/light cycle. Following headbar mounting the mice were single-housed since they tend to nibble on the 3D-printed plastic headbar.

### Headbar mounting

Mice were anesthetized within an induction box under isoflurane, the head was shaved and analgesics were given (Buprenorfine 0.1mg/Kg, Carprofene 5mg/Kg, I.P.). A few minutes later the animal was mounted within a custom-made head-holder which freely tilting the head in any direction (Haidarliu, 1996; Slotnick, 1972), and a headbar was mounted stereotactically as specified in the main text.

### Histology

For histological reconstructions the tip of glass patch pipette was dipped within melted DiI (Merck). The pipette was directed through a craniotomy into the coordinates of the amygdala (from bregma in mm: 1.7 posterior, 3.0 lateral, 3.8-4.7 ventral), the mammillothalamic tract (2.0 posterior, 0.63 lateral, 4.6 ventral) or the subthalamic nucleus (2.0 posterior, 1.7 lateral, 4.45 ventral) and was left there for 15 minutes. At the end of the experiment, the mice were over-anesthetized and perfused transcardially with 2.5% paraformaldehyde, and their brains were removed and postfixed for 24h in the perfusion solution. Brains were then immersed in PBS solution with additional 30% sucrose for 24h and then cut in a microtome (80 μm thick, SM 2000R; Leica, Heidelberg, Germany).

Brain slices were mounted on slides and scanned using ZEN software (Zeiss) by a confocal microscope (LSM-880, Zeiss). Fluorescence signals of YFP and Dil were acquired with compatible filter sets (ex 470/40, em 525/50 for YFP, ex 545/25, em 605/70 for DiI). Images were stitched and exported using ImageJ software.

### Histological verification of electrode position

Electrodes carrying DiI were positioned according to the coordinates of the amygdala, MT or STh. Using images from the confocal microscope (Fig. 3), we measured the distance between the electrode and the center of the amygdala. The distance of the electrode from the midline and the center of the amygdala were measured only in the mediolateral axis since the thickness of the slices does not allow to faithfully estimate the position within the amygdala and because the elongated shape of the amygdala makes it difficult to estimate the actual rostrocaudal position. The distances were measured semi-automatically by a custom-made software in LabVIEW in which the user marked the midline and the DiI position. Sometimes due to histological processes the tissue can shrink or expand (Hillman and Deutsch, 1978). However, based on measured distances between the STh marks (which are supposed 3.4 mm apart), we estimated that the size of the slices was changed less than 5%, with the error being positive in half the mice and negative in the other half. The lack of consistency in sign indicates the position measurements did not suffer from a systematic error, so they were not corrected.

### Electrophysiology and data acquisition

Electrophysiological signals were acquired using an Axoclamp-700B amplifier (Molecular Devices), lowpassed at 3 kHz before being digitized at 10 kHz (PCI-6221, National Instrument) by a custom software (Labview; RRID:SCR_014325). Data was processed, analyzed and presented using MATLAB (Mathworks, RRID:SCR_001622). Borosilicate micropipettes (BF150-86-10, Sutter instruments, Novato, CA) were pulled (P-97, Sutter instruments) to produce juxtacellular and intracellular electrodes carried by the optopatcher. The Optopatcher (#663843, A-M systems, WA) was used to optogenetically activate ChR2 expressing neurons with blue light (MDL-III-450L/ 1∼80mW, CNI lasers, P.R.China)

### Juxtacellular recording

Following a recovery period of at least three days from headbar implantation, the animals were anesthetized (∼1.5% isoflurane) and mounted on the DeepTarget, where a craniotomy (1 mm radius) over the amygdala was performed. Juxtacellular pipettes (12–18 MΩ) filled with artificial cerebrospinal fluid (containing in mM: 124 NaCl, 26 NaHCO3, 3 KCl, 1.24 KH2PO4, 1.3 MgSO4, and 2.4 CaCl2) were lowered to 3.8 mm beneath the dura and by monitoring changes in electrode resistance we looked for cells up to 4.7 mm that responded to light stimulation.

### Awake intracellular recording

Following a recovery period of at least three days from headbar implantation, animals were anesthetized (∼1.5% isoflurane), mounted on the DeepTarget and injected with carprofen (5 mg/Kg). Under anesthesia a craniotomy (1 mm radius) was performed over the amygdala.

The craniotomy was covered with warm agar and sealed with silicone elastomer (Body Double™ Fast Set, Smooth-On, Inc., Macungie, PA). Following the procedure the animals were returned to their home-cage to recover for 2 hours. After recovery animals were anesthetized, mounted back onto the DeepTarget while the silicone elastomer was removed, and then the anesthesia was disconnected and the recording session started. In order to perform intracellular recordings we used patch pipettes (6–9 MΩ) filled with an intracellular solution (containing in mM: 136 K-gluconate, 10 KCl, 5 NaCl, 10 HEPES, 1 MgATP, 0.3 NaGTP, and 10 phosphocreatine, 310 mOsm) we used the same procedures described in the ‘Juxtacellular recording’ section. In this case we tried to form a seal and get a whole-cell patch clamp configuration. Typically successful recordings were accomplished only by pipettes subsequent to the first, since the first one penetrated through the dura and cleared the way down to the amygdala.

## Data, Software and Designs Availability

Headbars and system designs are available at: https://drive.google.com/drive/folders/1AqFj6AOIdejWr7ALgYZZJnX_I7lNAdxw?usp=sharing

Other data are available on request. Please contact the lead Y.K‥

## Acknowledgements

YK is incumbent of the Marianne Manoville Beck Research Fellow Chair in Brain Research. We thank Rebekah Tumasus for the histological reconstructions. We thank Benny Pasmantirer and Georges Ankaoua from the Instruments Design Section in the Weizmann Institute of Science for designing the systems. This work was supported by grants from the DFG-SFB 1089, 01EW1606 - DeCipher EraNet Neuron, HFSP, Israel Science Foundation (ISF 1539/17) and Minerva, as well as by a research grant from the Marianne Manoville Beck Laboratory for Research in Neurobiology in Honor of her Parents Elisabeth and Miksa Manoville.

## Author contributions

Conceptualization, Y.K.; Methodology, Y.K, M.S, I.L; Investigation, Y.K.; Writing – Review & Editing,Y.K, M.S, I.L; Funding Acquisition, I.L.

## Abbreviations

BLA: - Basolateral amygdala
ML: - Mediolateral
MT: - Mammillothalamic tract
STh: - Subthalamic nucleus

